# Monarch butterfly *Cryptochrome 1* loss-of-function mutants reveal differences in light entrainment of 24-hour behavioral rhythms in insects

**DOI:** 10.1101/2023.08.04.552044

**Authors:** Samantha E. Iiams, Guijun Wan, Jiwei Zhang, Aldrin B. Lugena, Ying Zhang, Ashley N. Hayden, Christine Merlin

**Author notes:** Corresponding author: Christine Merlin, Texas A&M University, Department of Biology, Biological Sciences Building East, 3258 TAMU, 77843, College Station, TX, USA. Equal contribution (Guijun Wan, Jiwei Zhang).

## Abstract

Light is one of the strongest cues for entrainment of circadian clocks in most organisms. Previous work in *Drosophila melanogaster* (*dm*) has shown that entrainment relies on both the visual system and the circadian, blue-light photoreceptor Cryptochrome (dmCRY). Here, we used the monarch butterfly *Danaus plexippus* (*dp*) to test conservation of this mechanism among insects and the relative importance of monarch *Cryptochrome 1* (*dpCry1*) in the entrainment of its clock *in vivo*. We showed that loss of functional *dpCry1* abolishes adult circadian eclosion behavior and molecular circadian rhythms in the monarch brain. These rhythms can be restored by entrainment to temperature cycles, demonstrating that the core circadian clock is intact in *dpCry1* mutants. Importantly, we showed that rhythmic flight activity is also disrupted in *dpCry1* mutants but not in the visually impaired *dpNinaB1* mutants, suggesting that unlike *Drosophila* light-entrainment of the monarch circadian clock relies solely on dpCRY1 photoreception.

## INTRODUCTION

CRYPTOCHROMES (CRYs) are an ancient class of blue-light and ultraviolet (UV)-sensitive flavoproteins present in a wide range of organisms [1-3] that evolved from photolyases responsible for the repair of UV-generated pyrimidine dimers [4, 5]. Although CRYs have retained the N-terminal photolyase-related (PHR) domain, which contains the chromophore access cavity for binding of the flavin adenine dinucleotide (FAD) cofactor, they have lost the DNA repair capability of a photolyase [1-3] and classically function in the circadian system. Two types of CRYs, type 1 CRYs and type 2 CRYs, can be differentiated based on their circadian function. Type 2 CRYs, present in mammals and all insects studied so far with the notable exception of *Drosophila melanogaster* (*dm*), function as circadian repressors of the CLOCK (CLK):BMAL1 heterodimeric circadian transcription factor complex to generate circadian rhythms [6-11]. In contrast, type 1 CRYs, including *Drosophila* dmCRY, are light-sensitive and appear to function in the entrainment of the core circadian clock to generate circadian rhythms [12-14]. In type 1 CRYs, the C-terminal region contains a group of tryptophan (Trp) residues essential for enabling the photosensing capabilities of CRY by reducing the FAD chromophore into its active state [15-19], likely driving conformational changes which allows CRY to initiate downstream signaling pathways [20-23].

*Drosophila* dmCRY has been firmly established as the primary blue-light circadian photoreceptor for resetting the circadian clock cell autonomously to the daily light:dark (LD) cycle. DmCRY is rapidly degraded after light exposure and mediates the degradation of one of its partners, TIMELESS (dmTIM), in a light-dependent manner [12, 24]. A point mutation altering a key residue for FAD binding, named *cry*^*baby*^ (*cry*^*b*^), leads to severe entrainment defects, including a loss of light-dependent dmTIM degradation and the inability to phase-shift circadian response to light pulses [12, 13]. Curiously, *cry*^*b*^ mutants and a fly *cry*^*0*^ full knockout were shown to still retain behavioral rhythms in adult emergence from their pupal case and locomotor activity, and were also entrainable to new LD regimes [13, 25], suggesting the existence of other CRY-independent photoreceptors and pathways for entraining *Drosophila* circadian behaviors. This was confirmed by rendering the fly clock completely blind after eliminating all photoreceptors with a double mutant for *cry*^*b*^ and *glass*, a transcription factor necessary for the development of photoreceptors in the eyes and Hofbauer-Buchner eyelets [26]. Further characterization has shown that in addition to dmCRY, signal transduction pathways utilizing phospholipases encoded by *no receptor potential A* (*norpA*) and *phospholipase C* (*Plc21C*), as well as several rhodopsin (Rh) photoreceptors, Rh1, Rh5, Rh6, and Rh7 acting either in the eyes or the brain, play a role in *Drosophila* clock entrainment [27-30]. Whether such a complex network is also required for circadian clock entrainment in other insect species remains unknown.

To date, the role of light-sensitive CRY in clock entrainment in other insects has been limited to transgenic expression of mosquito CRY1s in *Drosophila* [31] and to *in vitro* studies in a monarch butterfly, *Danaus plexippus* (*dp*), embryo-derived DpN1 cell line that possesses a light-driven clock [10, 14, 24]. Yet, the extent of CRY1’s contribution to *in vivo* eclosion and activity rhythms remains unexplored in species outside *Drosophila*. Here, using a previously generated *dpCry1* loss-of-function monarch knockout, we genetically demonstrated that dpCRY1 functions as the major photoreceptor in the entrainment of monarch daily and circadian rhythms. Loss of functional *dpCry1* abolished circadian adult eclosion behavior and molecular circadian rhythms in the monarch brain. We showed that circadian eclosion rhythms deficiency can be restored by entrainment to temperature cycles in *dpCry1* mutants but not in the core clock component *dpClk* null mutants, demonstrating that the core circadian clock is intact in *dpCry1* mutants and the mutation only disrupts entrainment to light. Importantly, we showed that rhythmic flight activity was also disrupted in *dpCry1* mutants but not in visually impaired *dpNinaB1* mutants, suggesting that, unlike *Drosophila*, light-entrainment of the monarch behavioral rhythms relies primarily on dpCRY1 photoreception and not on a complex interplay with other photoreceptors.

## RESULTS

### *DpCry1* loss-of-function monarch mutants exhibit arrhythmic circadian adult eclosion behavior and molecular rhythms

To investigate whether the role of light-sensitive CRY in circadian behavioral rhythms entrainment is conserved among insect species *in vivo*, we started by testing if a *dpCry1* loss-of-function mutation, a 2-bp deletion in the fourth exon that led to no detectable production of the corresponding truncated protein that would otherwise lack the Trp-containing C-terminal domain [32], would affect monarch circadian eclosion rhythms. Eclosion, the emergence of the fully mature adult from its chrysalis, is currently the only known behavior in monarchs that is both under circadian control and easily tractable in laboratory conditions via recording with infrared cameras [13, 33]. Monarchs *dpCry1* wild-type (*dpCry1*^*+/+*^) and homozygous knockout (*dpCry1*^*-/-*^) siblings were raised from newly oviposited eggs to day 8 of the pupal stage in 12 h:12 h LD at 25°C and then released into constant darkness (DD) one to two days prior to their expected eclosion. As expected, *dpCry1*^*+/+*^ monarchs exhibited circadian eclosion rhythms with a peak in the number of monarchs emerging around the anticipated timing of lights on (CT0), as previously described [7, 11, 33, 34] (Fig. 1 **A**). In contrast, adult *dpCry1*^*-/-*^ eclosed arrhythmically throughout the 24 h circadian day (Fig. 1 **A**). This result stands in sharp contrast with findings in *cry*^*0*^ null mutant flies, which exhibit circadian pupal eclosion similar to that of wild-type flies [25]. Quantification by qPCR of mRNA levels of two core clock genes, *period* (*dpPer*) and *timeless* (*dpTim*), in monarch brains over the circadian day revealed that both genes cycled robustly in brains of wild-type monarchs but were expressed arrhythmically and at constitutively high levels in brains of *dpCry1*^*-/-*^ (Fig. 1 **B**).

**Figure 1:**
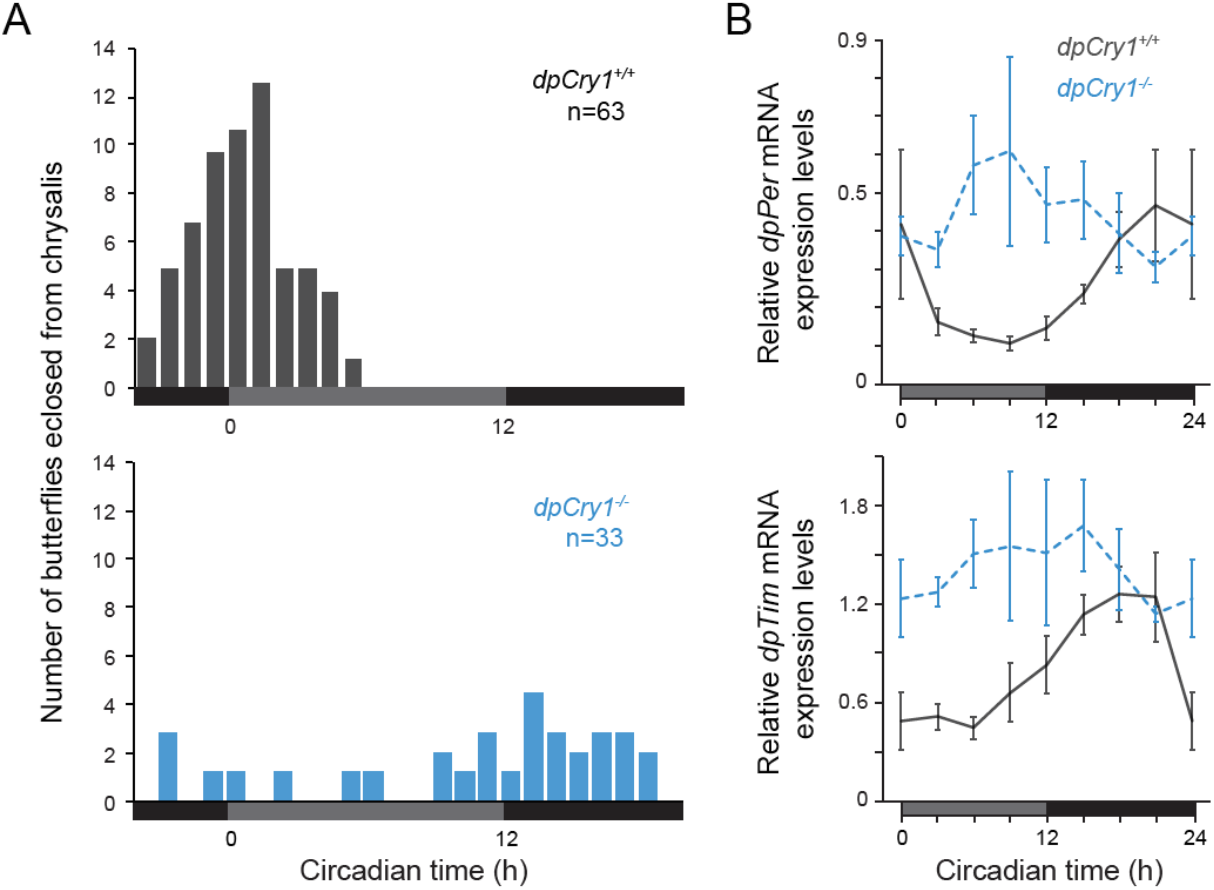
Circadian rhythms in eclosion behavior and brain mRNA clock genes levels are disrupted in monarch butterfly *dpCry1* knockouts. (**A**) Profiles of adult eclosion in constant darkness (DD) of wild-type (^+/+^; dark gray) and *dpCry1* homozygous mutant siblings (^-/-^; blue) entrained to 12 h:12 h light/dark (LD) cycles at 25°C throughout the larval stages and first week of pupation. Data from DD1 and DD2 are pooled and binned in 1 h intervals. Horizontal gray bars: subjective day; black bars: subjective night. Kolmogorov–Smirnov (KS) test maximum difference between *dpCry1*^*+/+*^ and *dpCry1*^*-/-*^: *p*= 0.00. KS normal distribution test: *dpCry1*^*+/+*^, *p*=0.23; *dpCry1*^*-/-*^, *p*=0.04. One-way ANOVA, *dpCry1*^*+/+*^ vs. *dpCry1*^*-/-*^, *p*=0.00. (**B**) Circadian expression of *period* (*dpPer*) and *timeless* (*dpTim*) in the brain of adult monarchs eclosed from **A** and re-entrained for at least 7 days to 12 h:12 h LD cycles at 25°C. *DpCry1*^*+/+*^: solid dark gray lines; *dpCry1*^*-/-*^: dashed blue lines. Values are mean ± SEM of four brains for each timepoint. One-way ANOVAs: *dpPer* in *dpCry1*^*+/+*^, *p*= 0.048; *dpPer* in *dpCry1*^*-/-*^, *p*=0.613; *dpTim* in *dpCry1*^*+/+*^, *p*=0.003; *dpTim* in *dpCry1*^*-/-*^, *p*=0.886. Two-way ANOVAs, interaction genotype × time: *dpPer, p*=0.036; *dpTim, p*=0.235.

### Temperature cycles re-entrain eclosion and molecular rhythms in *dpCry1* knockouts

To determine whether arrhythmic eclosion behavior and brain molecular rhythms observed in *dpCry1*^*-/-*^ mutant monarchs resulted from a deficiency in clock entrainment by light or in core clock function, we sought to bypass light by re-entraining the mutants to temperature cycles (TC), as TC has been shown to also act as a potent cue for entrainment in flies [13]. Monarchs were raised in 12 h:12 h LD at 25°C from eggs to the end of the fourth larval instar stage and subjected to constant light (LL) conditions for the duration of the fifth larval instar to eliminate any core clock oscillations from larval entrainment to light before entraining pupae to TCs [33] (Fig. 2 **A**). On the day of pupation, pupae were moved to a 12 h-15°C: 12 h-25°C TC in DD, in anti-phase to the LD cycle under which larvae were previously entrained, *i*.*e*. with low temperature coinciding to the previous light phase and high temperatures to the previous dark phase, until eclosion (Fig. 2 **A**). We found that both *dpCry1*^*+/+*^ wild-type and *dpCry1*^*-/-*^ mutant monarchs eclosed rhythmically with a strong peak at the beginning of the warm phase after the temperature rose from 15°C to 25°C, suggesting that eclosion behavior can be re-entrained to TC even in the absence of a functional dpCRY1 (Fig. 2 **B**, *Left*). To exclude the possibility that the observed TC entrainment was not caused by an acute response to the increase in temperature but was indeed under circadian clock control, the same experiment was performed using loss-of-function mutants of a core clock gene activator, *clock* (*dpClk*) [34], and their wild-type siblings. In contrast to wild-type monarchs, which eclosed rhythmically within the first two hours of the warm phase, *dpClk*^*-/-*^ eclosed randomly throughout it, demonstrating that a functional circadian clock is necessary for the entrainment of monarch eclosion to TCs (Fig. 2 **B**, *Right*). To test if re-entrainment of behavioral rhythms by TC would be mirrored at the molecular level, we quantified *dpPer* and *dpTim* mRNA levels over the 12 h-15°C: 12 h-25°C TC in DD in the brain of both adult wild-type and *dpCry1*^*-/-*^ mutants. We found that *dpPer* and *dpTim* mRNA rhythms were robustly entrained by TC in the brains of wild-type monarchs, with peaks of expression at the transition from warm to cold phases and troughs in the early warm phase (Fig. 2 **C**). Importantly, a similar trend was observed in the brains of *dpCry1*^*-/-*^ mutants, where *dpTim* exhibited rhythms indistinguishable from those in wild-type brains and *dpPer* also cycled albeit with a lower amplitude than in wild-type (Fig. 2 **C**). Together, these results demonstrate that *dpCry1*^*-/-*^ monarchs possess an intact core circadian clock that can be synchronized to TC. Thus, the unexpected loss of eclosion rhythms under LD conditions in *dpCry1*^*-/-*^ mutants suggests the intriguing possibility that, unlike in *Drosophila*, dpCRY1 may function as the sole light-entrainment pathway of circadian rhythms in monarchs.

**Figure 2:**
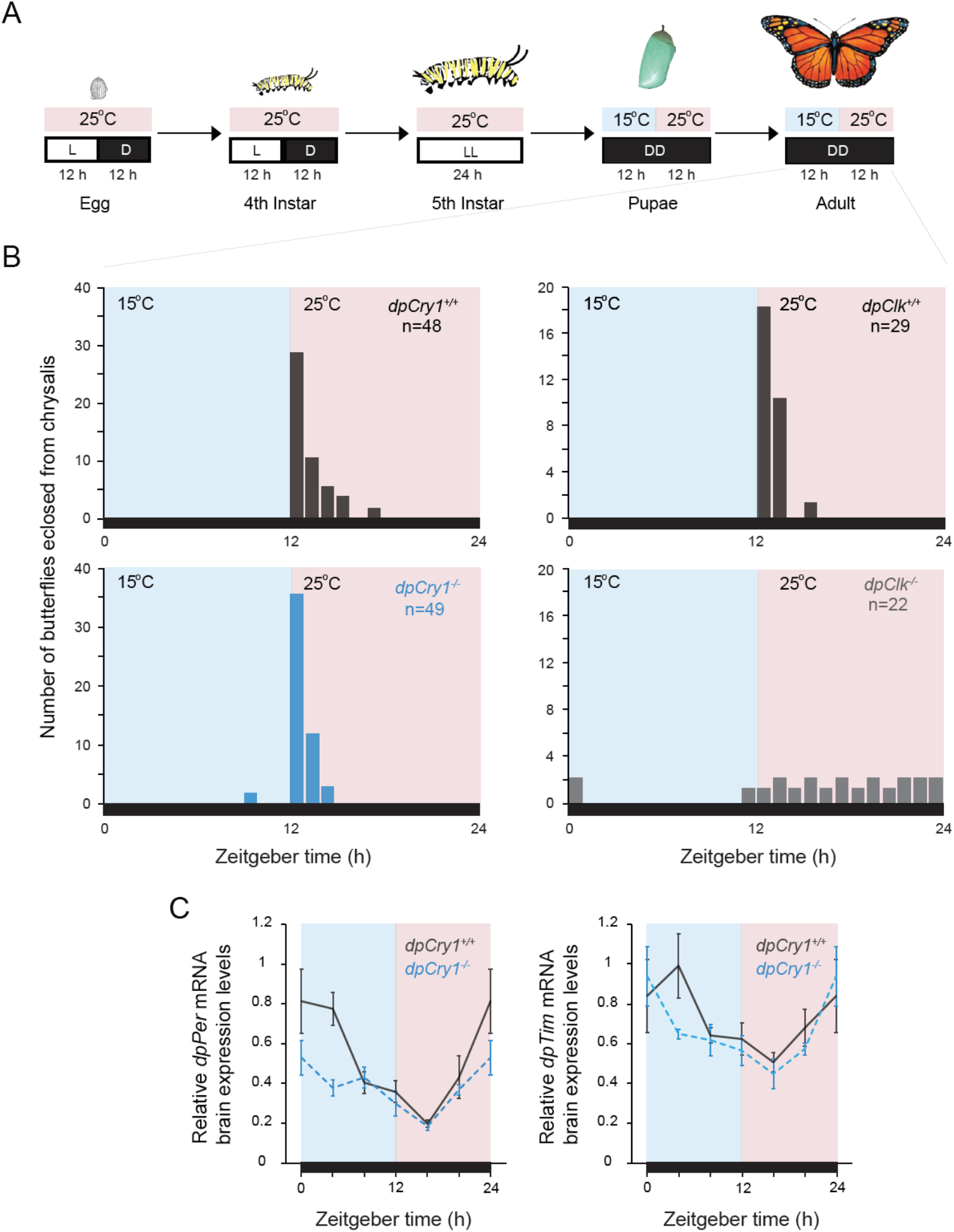
Temperature cycles re-entrain rhythmic eclosion behavior and brain molecular rhythms in *dpCry1* knockouts. (**A**) Schematic of entrainment by LD and temperature cycles at different developmental stages from egg to adult. Monarchs were raised under 12:12 LD cycles at 25°C from egg to fourth instar larvae, transferred into constant light (LL) during the fifth instar, and then placed in constant darkness (DD) in temperature cycles at 15°C for 12 h and 25°C for 12 h throughout pupal and adult stages. (**B**) Profiles of adult eclosion after pupal entrainment for ∼10 days to 12 h-15°C/ 12 h-25°C cycles. Black horizontal bars: constant darkness. (*Left*) Profiles of *dpCry1*^*+/+*^(*top*, dark gray) and *dpCry1*^*-/-*^ siblings (*bottom*, blue). KS test maximum difference between genotypes: *p*= 0.639. KS normal distribution test: *dpCry1*^*+/+*^, *p*=0.00; *dpCry1*^*-/-*^, *p*=0.00. One-way ANOVA between *dpCry1*^*+/+*^ and *dpCry1*^*-/-*^: *p*=0.016. (*Right*) Profiles of *dpClk*^*+/+*^ (*top*, dark gray) and *dpClk*^*-/-*^ siblings (*bottom*, light gray). KS test maximum difference between genotypes: *p*= 0.00. KS normal distribution test: *dpClk*^*+/+*^, *p*=0.00; *dpClk*^*-/-*^, *p*=0.24. One-way ANOVA between *dpClk*^*+/+*^ and *dpClk*^*-/-*^: *p*=0.005. (**C**) Expression of *dpPer* and *dpTim* in the brain of adult wild-type and *dpCry1* knockouts monarchs eclosed from **B** after at least 7 days of adult entrainment to 12 h-15°C/ 12 h-25°C in DD. *DpCry1*^*+/+*^, solid dark gray lines; *dpCry1*^*-/-*^, dashed blue lines. Values are mean ± SEM of four brains for each timepoint. One-way ANOVAs: *dpPer* in *dpCry1*^*+/+*^, *p*= 0.0007; *dpPer* in *dpCry1*^-/-^, *p*=0.006; *dpTim* in *dpCry1*^*+/+*^, *p*=0.09; *dpTim* in *dpCry1*^-/-^, *p*=0.012. Two-way ANOVAs, interaction genotype × time: *dpPer, p*=0.045; *dpTim, p*=0.394.

### Daytime patterns of adult flight activity are disrupted in *dpCry1* knockouts but not in blind *dpNinaB1* knockouts

Because the visual system and CRY-independent light entrainment pathways may not be fully developed until after adult emergence in monarchs, we sought to test the relative contribution of dpCRY1 on locomotor rhythms in adults. To this end, we used custom-built flight mills to assess daily patterns in flight activity of adult *dpCry1*^*-/-*^ mutants and wild-type siblings. Individual monarchs were tethered and suspended to one end of a balanced mill arm placed inside a plastic barrel allowing them to freely fly on a horizontal, circular plane, in 15 h: 9 h LD conditions at 21°C. Activity was recorded over 3 days using an infrared beam placed above the butterfly’s rotational flight path, and the distance flown was calculated based on the number of revolutions per 1 h time-bins. LD conditions were chosen over DD as monarch flight activity is inhibited in the absence of light (not shown), impeding formal tests of circadian activity.

Despite this limitation, we observed a significant impact of loss of functional *dpCry1* on rhythmic flight behavior in LD. In contrast to wild-type monarchs, which flew in a rhythmic pattern within the 15 h light phase with a peak in the middle of the day, *dpCry1*^*-/-*^ flew on average equal amounts throughout the light phase over the 3 days of recordings (Fig. 3 **A, B**). These results strongly suggest that *dpCry1* loss-of-function disrupts adult rhythmic flight activity, similar to its effect on circadian eclosion behavior.

**Figure 3:**
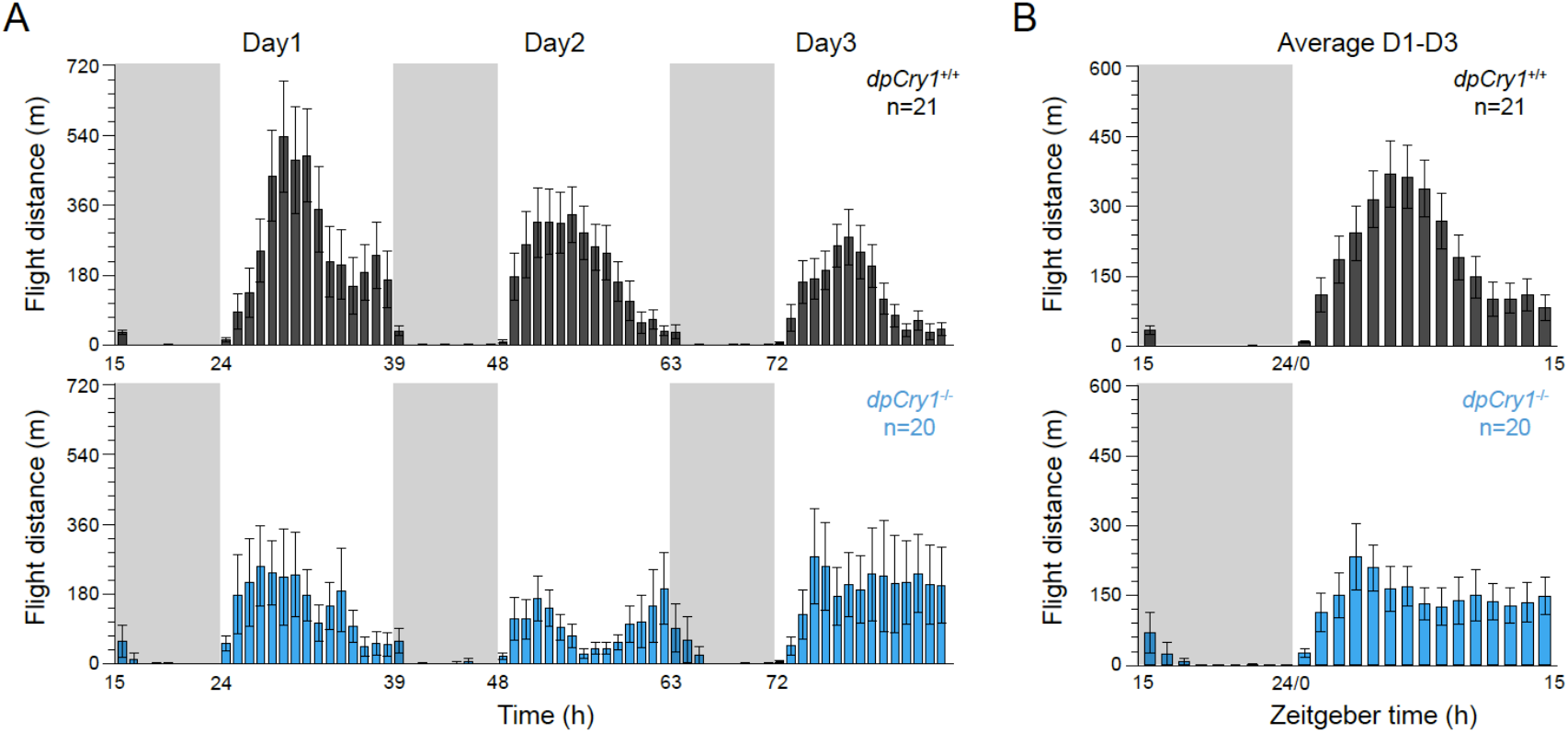
Rhythmic flight activity exhibited by wild-type monarchs during the light phase is abolished in *dpCry1* knockouts. (**A**) Profiles of flight activity in 15 h:9 h LD cycles over three consecutive days measured as distance flown in a flight mill by *dpCry1*^*+/+*^ (dark gray, *top*) and *dpCry1*^-/-^ siblings (blue, *bottom*). Data are binned in 1 h intervals. Gray shading: night; no shading: day. (**B**) Averaged flight activity over the three days of recording. Legends are as in (**A**). Meta2d rhythmicity analysis for the 15 h daytime flight period: *dpCry1*^*+/+*^, *p=*0; *dpCry1*^-/-^, *p=*1.

We also tested whether other light-sensing pathways could contribute to the daily rhythmic flight activity of monarchs. To determine the possible contribution of photoreceptors in the compound eyes, we blocked the light input to the eyes of wild-type and *dpCry1*^*-/-*^ mutant monarchs by painting them with black enamel paint [35, 36]. We found that flight activity patterns of wild-type monarchs with black painted eyes were indistinguishable from those of unpainted wild-type, with rhythmic flight activity restricted to the light phase and peaking in the middle of the day (Fig. S1, **A, B**). *DpCry1*^*-/-*^ monarchs with black painted eyes also showed a disrupted rhythm similar to that of unpainted *dpCry1*^*-/-*^ (Fig. S1, **A, B**). These results strongly supported the ideas that dpCRY1 acts as a deep-brain photoreceptor for the entrainment of behavioral rhythms and that eye photoreceptors do not play a role in clock entrainment in the monarch. Another key light-sensing pathway in *Drosophila* circadian photoentrainment has recently been shown to rely on the deep-brain rhodopsin 7 (Rh7) [30]. Although single mutants for either fly *dmCry* or *dmRh7* only exhibited minor defects on photoentrainment, the double mutant was impaired with increased period length in locomotor activity and percentage of flies exhibiting arrhythmicity. To test whether a deep-brain opsin could also contribute to light-entrainment of activity rhythms in monarchs, we tested the flight activity of a *dpNinaB1* loss of function mutant. In this mutant, the production of retinal, the chromophore that renders opsins light-sensitive, is impaired in both eyes and brain resulting in individuals that are blind, including to seasonally changing photoperiods [35]. We found that both wild-type and *dpNinaB1*^*-/-*^ mutant monarchs exhibited daylight rhythms in flight activity with the expected peak during midday (Fig. 4 **A, B**; *top two panels*). We also observed a notable increase in nighttime flight activity in the *dpNinaB1*^*-/-*^ mutants. To then test if loss of function of both dpCRY1 and dpNINAB1 would have additive or synergistic effects on activity rhythms, we recorded daily flight activity of double mutants. As compared to rhythmic controls, *dpCry1*^*-/-*^*;dpNinaB1*^*-/-*^ lost rhythms in daylight flight activity and exhibited a lack of repression of flight at the onset of darkness, suggesting that day flight and night flight may be differentially regulated by separate photoreceptor pathways (Fig. 4 **A, B**; *bottom two panels*). Furthermore, the increase in nighttime activity observed in both *dpNinaB1*^*-/-*^ and *dpCry1*^*-/-*^*;dpNinaB1*^*-/-*^ mutants compared to controls raises the intriguing possibility that the vitamin A pathway and/or an opsin is involved in repressing activity at night or in promoting sleep (Fig. 4 **A, B**).

**Figure 4:**
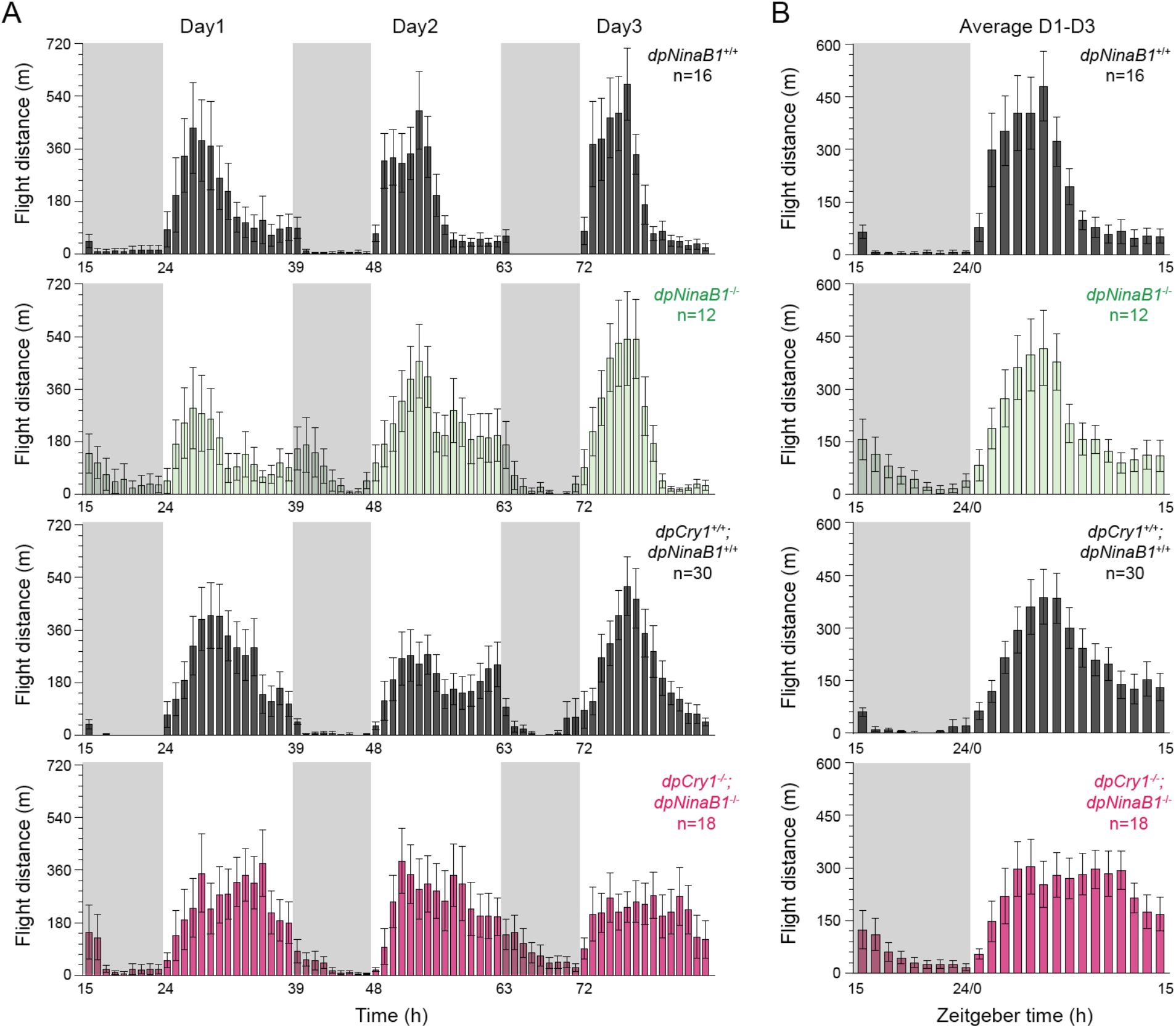
Genetic loss of opsin-based photoreception does not affect daytime rhythmic flight activity of monarchs. (**A**) Profiles of flight activity in 15 h:9 h LD cycles over three consecutive days measured as distance flown in a flight mill by blind *dpNinaB1*^*-/-*^(green), *dpCry1*^*-/-*^*;dpNinaB1*^*-/-*^ double homozygous knockouts (pink), and their corresponding wild-type siblings (*dpNinaB1*^*+/+*^ and *dpCry1*^+/+^*;dpNinaB1*^*+/+*^ respectively, black). Data are binned in 1 h intervals. Gray shading: night; no shading: day. (**B**) Averaged flight activity over the three days of recording. Legends are as in (**A**). Meta2d rhythmicity analysis for the 15 h daytime flight period: *dpNinaB1*^*+/+*^, *p*=0; *dpNinaB1*^*-/-*^, *p<*0.0001; *dpCry1*^*+/+*^; *dpNinaB1*^*+/+*^, *p=*0; *dpCry1*^*-/-*^; *dpNinaB1*^*-/-*^, *p=*0.002.

## DISCUSSION

Over the past 15 years, comparative analyses of clock genes in insects have revealed a greater diversity of circadian clock molecular make-ups within insects than that of the pioneering insect model *Drosophila melanogaster* [9, 37-39]. A distinctive feature that has attracted much attention was the presence of a mammalian-like type 2 CRY in non-Drosophilid insect species that functions as a circadian transcriptional repressor [7, 9, 11]. Less attention has been paid to the diversity of light-entrainment pathways. While type 1 CRY has been lost in some insect species, including the honeybee *Apis mellifera* and the flour beetle *Tribolium castaneum*, it is found in others like moths, butterflies, and mosquitos [9, 40, 41]). Whether type 1 CRYs function as circadian photoreceptors in concert with other signaling pathways for the light-entrainment of daily and circadian behaviors in non-Drosophilid species remains unknown, as our current knowledge has remained limited to *Drosophila melanogaster* [12, 13, 25-30].

To our surprise, characterizing the relative importance of the ortholog of dmCRY in the entrainment of 24 h rhythms in the monarch butterfly revealed a fundamental difference in the complexity of the photoreceptive network for the *in vivo* light-entrainment of the circadian clock in insects. Unlike *Drosophila cry* null mutants, in which behavioral rhythmicity persists due to additional dmCRY-independent rhodopsin photoreceptors and signal transduction pathways for light-entrainment of the circadian clock [27, 28, 30], monarch *dpCry1* knockouts eclosed arrhythmically even after long-term entrainment to LD conditions. The rescue of rhythmic eclosion and molecular rhythms of *dpPer* and *dpTim* in adult brains by TC entrainment in this mutant demonstrated that its core circadian clock machinery was intact and functional, confirming a light-entrainment specific deficiency in *dpCry1* knockouts. Although eclosion behavior was tightly linked to the increase in temperature in the *dpCry1* mutant entrained to TC, molecular rhythms of *dpPer* and *dpTim* exhibited anticipatory phases of decreasing expression starting in the cold and continuing well into the warm phase before increasing again, arguing against a masking effect of rising temperatures in eclosion behavior. The ability to entrain circadian clocks to TCs has been documented both in insects [13, 42, 43] and mammals [44, 45], and has been found to be mediated by nocte and the Ionotropic Receptor 25a (IR25a) in *Drosophila* [46, 47]. Orthologues of these temperature signal mediators are present in the monarch genome (data not shown). Future experiments knocking out these genes via CRISPR/Cas9 could establish if they too serve a similar role in temperature entrainment in the monarch.

Although loss of circadian rhythms in eclosion in *dpCry1* mutants after LD entrainment supports the idea that dpCRY1 functions as the primary photoreceptor for entrainment of the monarch clock to light *in vivo*, we have not excluded the formal possibility that loss of rhythmic eclosion behavior in these mutants could result from dpCRY1-independent pathways not yet functional in the monarch pupal brain. In *Drosophila, cry* null mutants not only maintained rhythms in adult activity, but also eclosed with a ∼24 h rhythm, suggesting that CRY-independent pathways are functional by the time of adult eclosion, at least in this species [25]. In support of this hypothesis, developmental studies of the fly visual system have revealed that the optic nerve connects to the circadian clock early in the embryonic stage and that *Rh5, Rh6*, and *norpA* are already expressed within larval retina cells [48]. Whether such pathways are also formed in the larval stage of monarch butterflies remains unknown. To exclude the possibility that loss of eclosion circadian rhythms in *dpCry1* mutants could be due to yet undeveloped CRY-independent, opsin-based, light sensing pathways at the pupal stage, we established a new behavioral assay to assess 24 h rhythms in adult flight activity. As previously observed anecdotally, monarchs remain almost completely inactive at night. However, flight activity during the light phase was itself rhythmic with a peak near mid-day in wild-type monarchs, whereas such daytime rhythms were abolished in the *dpCry1* mutants.

Surprisingly, blocking light input to the compound eyes with black paint did not abolish wild-type rhythms nor affected the phase of the peak of activity, suggesting that CRY-independent light sensing pathways in the compound eye, such as those utilized in *Drosophila*, are not playing a significant role in the entrainment of daytime rhythmic flight activity in monarchs. Furthermore, our data demonstrate that a loss-of-function mutation in *dpNinaB1*, the gene encoding the rate-limiting enzyme in the vitamin A pathway responsible for the conversion of β-carotene into retinal, does not impact daytime patterns of flight activity. Because these mutants are blind and lack a functional vitamin A pathway in the brain [35], our data indicate that the contribution of brain retinal-opsins in the regulation of daytime rhythms in flight activity can also be excluded and highlight dpCRY1 as the primary photoreceptor for light-entrainment of monarch flight activity rhythms. However, in contrast to wild-type monarchs that become inactive almost immediately upon lights off, *dpNinaB1* mutants continued to exhibit flight activity well into the night. This unexpected finding may suggest that a retinal-opsin plays a role in regulating the repression of monarch flight activity in darkness. One possible mechanistic explanation for this mutant phenotype is that the lack of retinal could leave the relevant opsin in a constitutively active state, as observed in vertebrates [49, 50], prolonging flight activity into the night. Identifying the opsin(s) involved in repressing night-time monarch flight activity, among the ones present in the monarch genome, will be necessary to eventually test this hypothesis. Together, our data suggest the existence of two separate photoreceptor signaling pathways functioning in the monarch brain to drive rhythmic flight activity; one that relies on dpCRY1 to drive daytime rhythms in adult flight activity, and another that likely employs a retinal-opsin to sense the onset of darkness and repress flight activity in light:dark conditions.

The combined loss of eclosion rhythms, daytime adult flight activity rhythms, and molecular rhythms in *dpCry1* mutants highlights the primary role of dpCRY1 in the entrainment of the monarch activity rhythms *in vivo*, unlike *Drosophila* which relies on both dmCRY and rhodopsins/phospholipase-based signal transduction pathways. By uncovering significant species-specificity in the complexity (or lack thereof) of light-entrainment mechanisms between insect species, our work underscores a need for future mechanistic studies in a greater number of insect/invertebrate species to better understand the evolution of entrainment of circadian rhythms by light across invertebrates.

## LIMITATIONS OF THE STUDY

Adult monarch butterflies do not exhibit significant (and thus quantifiable) locomotor or flight activity in constant darkness, even after being entrained to light:dark regimens for at least seven days, an amount sufficient to entrain circadian rhythms in brain clocks. The lack of sustained flight activity rhythms in constant conditions thus precluded analyses of truly circadian flight activity rhythms. Future investigations leveraging our discovery that flight activity increases during the dark phase in the absence of a functional vitamin A pathway may prove useful in identifying conditions in which such studies will be possible.

## ACKNOWLEDGEMENTS

We thank Catherine Bogdan, Anna Subonj, Alec Judd, Alyssa Bennett and Jenna Coleman for their assistance with monarch rearing. The work was supported by NSF IOS-1456985, NSF IOS-1754725 and NSF IOS-2224154 to C.M.

## AUTHOR CONTRIBUTIONS

Author contributions: S.E.I. and C.M. designed the research; S.E.I., G.W., J.Z., A.B.L., Y.Z., A.N.H., and C.M. performed research; S.E.I., G.W., J.Z., A.B.L., and C.M. analyzed data; and S.E.I. and C.M. wrote the paper.

## STAR METHODS

### Resource availability

#### Lead contact

Further information and requests for resources and reagents should be directed to and will be fulfilled by the Lead Contact, Christine Merlin at.

#### Materials availability

This study did not generate new unique reagents.

### Experimental model and subject details

#### Mutant Monarch Butterfly Strains

Monarch *dpCry1* knockouts used throughout the experiments, *dpClk* knockouts used in the temperature cycle entrainment paradigm, and *dpNinaB1* knockouts used for flight activity assays were generated in prior studies via CRISPR/Cas9-mediated targeted mutagenesis [32, 34, 35]. *DpCry1;dpNinaB1* double knockouts and wild-type siblings were generated by crossing the *dpCry1* and *dpNinaB1* mutant strains to obtain and intercross double heterozygotes.

#### Monarch Butterfly Husbandry

For all experiments listed, monarch larvae were reared on a semi-artificial diet and adults were manually fed a 25% honey solution every day, as previously described [34].

For testing circadian eclosion behavior, wild-type and dp*Cry1* homozygous mutant monarchs were raised at 25°C with 70% humidity under the following light:dark cycle (LD) conditions: 15 h: 9 h LD from eggs to fifth instar larvae, 12 h: 12 h LD from days 1 to 8 of pupation, followed by one to two days of constant dark (DD). For quantification of gene expression in the adult brains, adults eclosed from this experiment were re-entrained to 12 h:12 h LD cycles for at least 7 days before brains were dissected under red light in the first day of DD, as previously described [35].

For testing eclosion behavior under temperature cycles in DD, wild-type, *dpCry1* and *dpClk* homozygous mutant monarchs were raised from eggs to fourth instar larvae in 12 h:12 h LD at 25°C with 70% humidity, transferred to constant light conditions (LL) at same temperature and humidity levels for the duration of the fifth instar, and then moved at pupation to DD in a 12 h-15°C/ 12 h-25°C temperature cycle in anti-phase to the previous larval LD cycle (*i*.*e*., 12 h of exposure to 25°C occurs in the larvae’s previous dark phase and 12 h of exposure to 15°C occurs in the larvae’s previous light phase). For quantification of gene expression in the adult brains, adults eclosed from this experiment were maintained in the same 12 h-15°C/ 12 h-25°C cycles in DD for at least 7 days before brain dissection.

For recordings of daily flight activity, monarchs were raised under 15-h: 9-h LD cycles at 25°C from eggs to adults and maintained under the same lighting conditions but at 21°C as adults for at least 7 days before flight activity recordings.

### Method details

#### Real-Time qPCR

For the circadian time course experiment, the brains of adult wild-type and *dpCry1* homozygous mutants were entrained to at least 7 days of 12 h: 12 h LD cycles and dissected during the first day of DD at circadian time (CT) CT0, CT3, CT6, CT9, CT12, CT15, CT18, and CT21 under red light in 0.5X RNA later (Invitrogen).

For the temperature entrainment time course experiment, brains of adult wild-type, *dpCry1* homozygous mutants were entrained to at least 7 days of 12 h-15°C/ 12 h-25°C in DD before being dissected at zeitgeber time (ZT) ZT0, ZT4, ZT8, ZT12, ZT16, and ZT20.

For both experiments, RNA extractions and gene expression quantifications were performed as previously described [35], using validated monarch *per, tim*, and control *rp49* primers [34]. The data were normalized to *rp49* as an internal control and normalized to the mean of one sample within a set for statistics.

#### Eclosion Behavior Assays

Adult eclosions were recorded inside an incubator using a video-tracking system, as previously described [7, 34]. For circadian eclosion assays, pupae were raised for 7-8 days in 12 h: 12 h LD cycles at 25°C and released in DD 1 to 2 days prior to emergence, as defined by the appearance of melanization on legs visible through the pupal case. Eclosion was observed on the first and second day of DD. For temperature cycle-entrained eclosion assays, eclosion was observed between 10-15 days after pupation while being continuously entrained to 12 h-15°C: 12 h-25°C in DD. The developmental delay in these conditions was caused by the overall temperature decrease. Eclosion data were analyzed and plotted as 1 h bins [34, 35].

#### Adult Flight Activity Assays

Rhythmic monarch flight activity was measured using custom-built flight mills individually placed inside plastic barrels, in which individual monarchs were suspended by a tether glued to their thorax on one end of the flight arm counterbalanced by a weight on the other side and allowed to freely fly in a circular horizontal plane, as previously described [32]. The number of revolutions of the arm at a time resolution of one minute was automatically recorded using an infrared beam connected to an automatic recording counter system. Each genotype of monarchs was tested blindly under a 15 h: 9 h LD cycle at 21°C. All monarchs were acclimated for at least 3 hours before the three days of motor activity tests and were fed daily with 25% honey solution ∼15 minutes at ZT12. Data were collected using a commercial data acquisition software (CHMBDD, MB-96) and the distance flown by each individual was calculated based on the number of revolutions per hour and the radius of the circular flight path, as previously described [32].

### Quantification and statistical analysis

#### Eclosion Behavior and Brain Molecular Rhythms

P-values were calculated using one-way ANOVAs or two-way ANOVAs followed by post-hoc analyses, and Kolmogorov–Smirnov tests using online calculators at https://www.icalcu.com/stat/anova-tukey-hsd-calculator.html, https://www.wessa.net/rwasp_Two%20Factor%20ANOVA.wasp [51], and http://www.physics.csbsju.edu/stats/KS-test.n.plot_form.html [52].

#### Adult Flight activity Rhythmicity

Rhythmicity analysis of monarch flight activities during the day was performed using MetaCycle [53] (Supplemental Table 1). For each genotype, the flight data taken for 3 consecutive days for each individual butterfly tested were considered as technical replicates, while all butterflies tested were treated as biological replicates. As flight activity is suppressed in darkness and only exhibits a rhythm during the light phase (from ZT0 to ZT15 in our experiments) peaking in the middle of the day, rhythmic analysis of the day flight activity data was restricted to ZT1 through ZT15. A meta2d analysis with default parameters except for the “minper” and “maxper”, which were both fixed at 15, was performed on the average of technical replicates for each butterfly. Due to uneven number of biological replicates between genotypes, rhythmicity tests were performed separately for each genotype.

## SUPPLEMENTAL INFORMATION

**Supplemental Figure 1:**
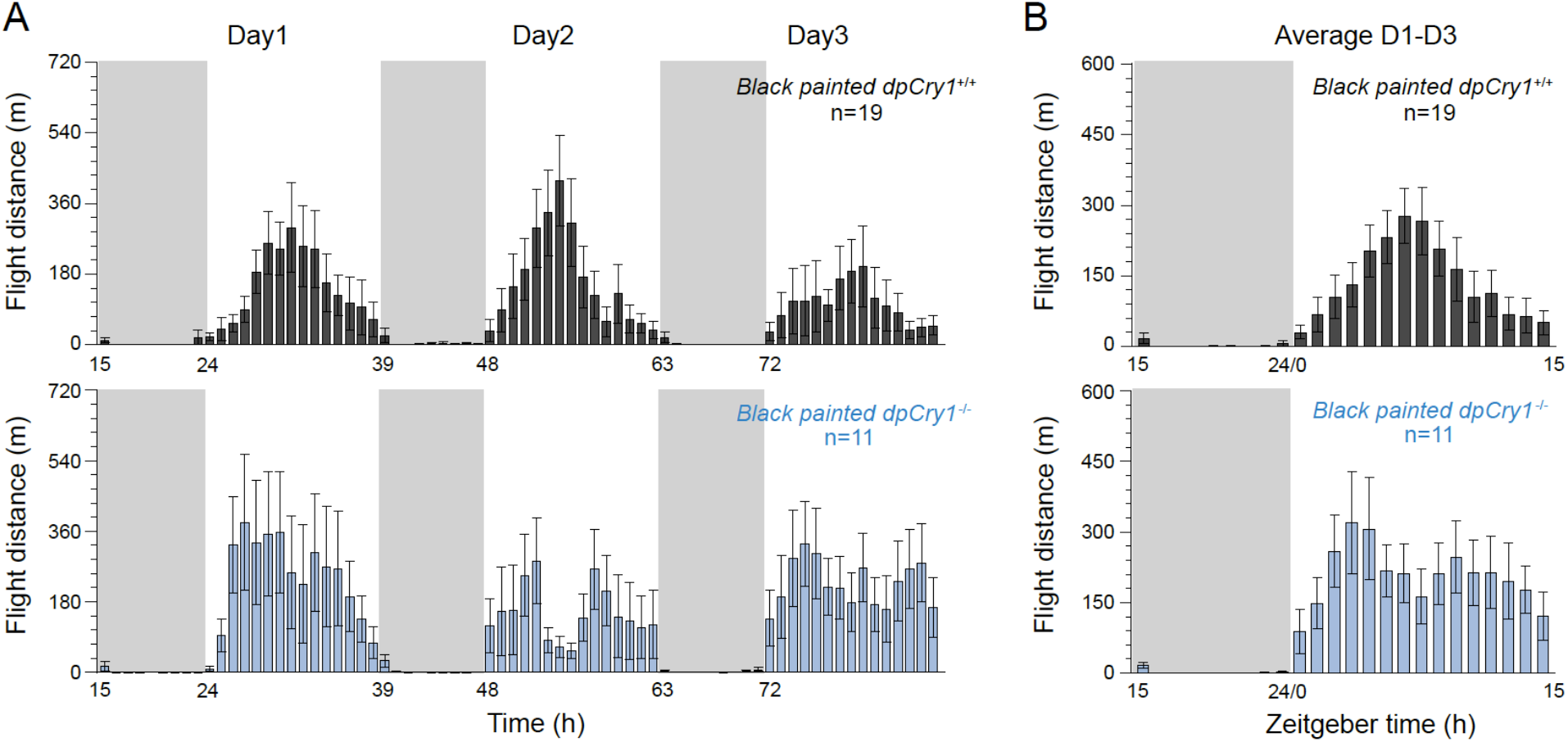
Blocking light-entrainment via the compound eyes does not affect rhythmic daytime flight activity. (**A**) Profiles of flight activity in 15 h:9 h LD cycles for three consecutive days measured as distance flown in a flight mill by *dpCry1*^*+/+*^ (dark gray, *top*) and *dpCry1*^*-/-*^ siblings (light blue, *bottom*) both with compound eyes painted black. Data are binned in 1 h intervals. Gray shading: night; no shading: day. (**B**) Averaged flight activity from the three days of recording. Legends are as in **A**. Meta2d rhythmicity analysis for the 15 h daytime flight period: *dpCry1*^*+/+*^, *p<*0.0001; *dpCry1*^-/-^, *p=*0.207.

**Supplemental Table 1:**
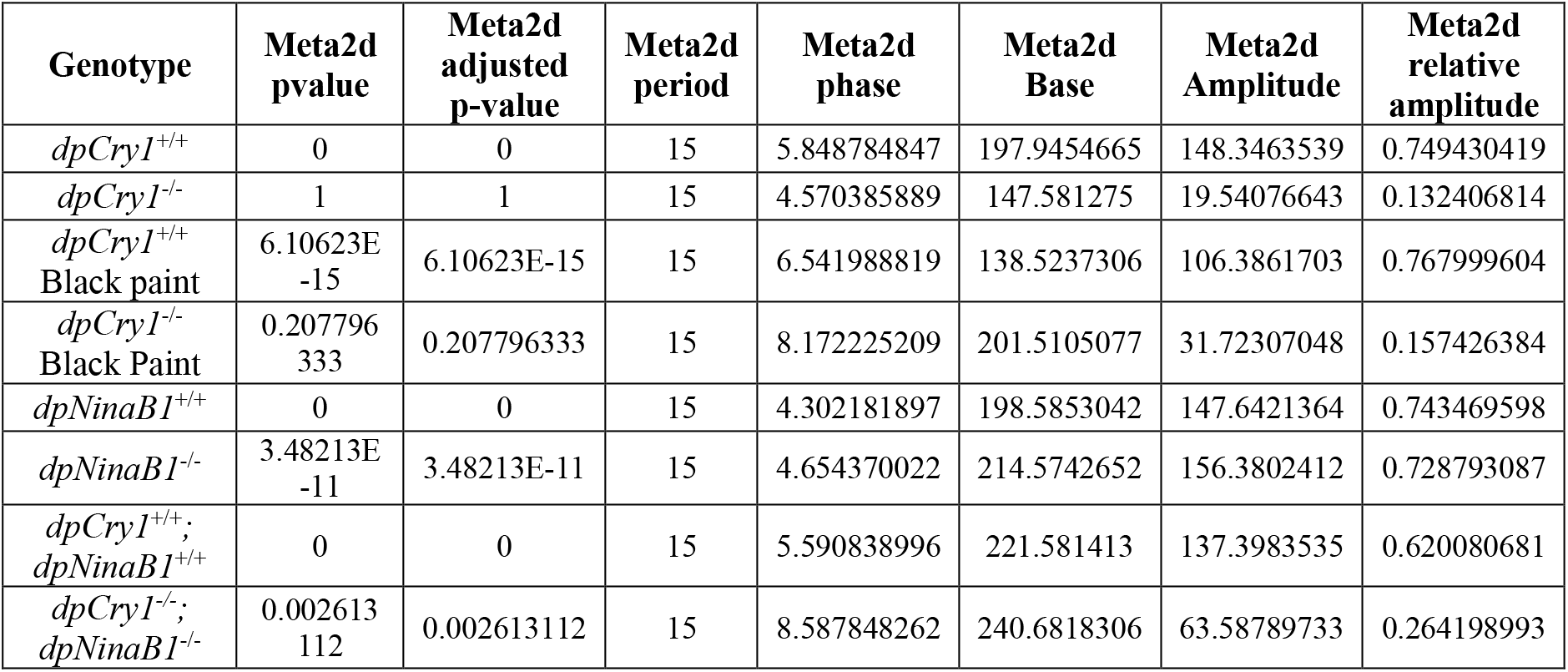
Rhythmicity analysis of flight data via MetaCycle.

## REFERENCES

1. Todo, T., Functional diversity of the DNA photolyase/blue light receptor family. Mutat Res, 1999. 434(2): p. 89–97.

2. Lin, C. and T. Todo, The cryptochromes. Genome Biol, 2005. 6(5): p. 220.

3. Mei, Q. and V. Dvornyk, Evolutionary History of the Photolyase/Cryptochrome Superfamily in Eukaryotes. PLoS One, 2015. 10(9): p. e0135940.

4. Michael, A.K., et al., Animal Cryptochromes: Divergent Roles in Light Perception, Circadian Timekeeping and Beyond. Photochem Photobiol, 2017. 93(1): p. 128–140.

5. Sancar, A., Structure and function of photolyase and in vivo enzymology: 50th anniversary. J Biol Chem, 2008. 283(47): p. 32153–7.

6. Kume, K., et al., mCRY1 and mCRY2 are essential components of the negative limb of the circadian clock feedback loop. Cell, 1999. 98(2): p. 193–205.

7. Merlin, C., et al., Efficient targeted mutagenesis in the monarch butterfly using zinc-finger nucleases. Genome Res, 2013. 23(1): p. 159–68.

8. van der Horst, G.T., et al., Mammalian Cry1 and Cry2 are essential for maintenance of circadian rhythms. Nature, 1999. 398(6728): p. 627–30.

9. Yuan, Q., et al., Insect cryptochromes: gene duplication and loss define diverse ways to construct insect circadian clocks. Mol Biol Evol, 2007. 24(4): p. 948–55.

10. Zhu, H.S., et al., The two CRYs of the butterfly. Current Biology, 2005. 15(23): p. R953–R954.

11. Zhang, Y., et al., Vertebrate-like CRYPTOCHROME 2 from monarch regulates circadian transcription via independent repression of CLOCK and BMAL1 activity. Proc Natl Acad Sci U S A, 2017. 114(36): p. E7516–E7525.

12. Emery, P., et al., CRY, a Drosophila clock and light-regulated cryptochrome, is a major contributor to circadian rhythm resetting and photosensitivity. Cell, 1998. 95(5): p. 669–79.

13. Stanewsky, R., et al., The cryb mutation identifies cryptochrome as a circadian photoreceptor in Drosophila. Cell, 1998. 95(5): p. 681–92.

14. Zhu, H.S., et al., Cryptochromes define a novel circadian clock mechanism in monarch butterflies that may underlie sun compass navigation. Plos Biology, 2008. 6(1): p. 138–155.

15. Chaves, I., et al., The cryptochromes: blue light photoreceptors in plants and animals. Annu Rev Plant Biol, 2011. 62: p. 335–64.

16. Kao, Y.T., et al., Ultrafast dynamics and anionic active states of the flavin cofactor in cryptochrome and photolyase. J Am Chem Soc, 2008. 130(24): p. 7695–701.

17. Lin, C., et al., Circadian clock activity of cryptochrome relies on tryptophan-mediated photoreduction. Proc Natl Acad Sci U S A, 2018. 115(15): p. 3822–3827.

18. Martin, R., et al., Ultrafast flavin photoreduction in an oxidized animal (6-4) photolyase through an unconventional tryptophan tetrad. Phys Chem Chem Phys, 2017. 19(36): p. 24493–24504.

19. Nohr, D., et al., Extended Electron-Transfer in Animal Cryptochromes Mediated by a Tetrad of Aromatic Amino Acids. Biophys J, 2016. 111(2): p. 301–311.

20. Berndt, A., et al., A novel photoreaction mechanism for the circadian blue light photoreceptor Drosophila cryptochrome. J Biol Chem, 2007. 282(17): p. 13011–21.

21. Ganguly, A., et al., Changes in active site histidine hydrogen bonding trigger cryptochrome activation. Proc Natl Acad Sci U S A, 2016. 113(36): p. 10073–8.

22. Hoang, N., et al., Human and Drosophila cryptochromes are light activated by flavin photoreduction in living cells. PLoS Biol, 2008. 6(7): p. e160.

23. Vaidya, A.T., et al., Flavin reduction activates Drosophila cryptochrome. Proc Natl Acad Sci U S A, 2013. 110(51): p. 20455–60.

24. Ceriani, M.F., et al., Light-dependent sequestration of TIMELESS by CRYPTOCHROME. Science, 1999. 285(5427): p. 553–6.

25. Dolezelova, E., D. Dolezel, and J.C. Hall, Rhythm defects caused by newly engineered null mutations in Drosophila’s cryptochrome gene. Genetics, 2007. 177(1): p. 329–45.

26. Helfrich-Forster, C., et al., The circadian clock of fruit flies is blind after elimination of all known photoreceptors. Neuron, 2001. 30(1): p. 249–61.

27. Ogueta, M., R.C. Hardie, and R. Stanewsky, Non-canonical Phototransduction Mediates Synchronization of the Drosophila melanogaster Circadian Clock and Retinal Light Responses. Curr Biol, 2018. 28(11): p. 1725–1735 e3.

28. Senthilan, P.R., et al., Role of Rhodopsins as Circadian Photoreceptors in the Drosophila melanogaster. Biology (Basel), 2019. 8(1).

29. Szular, J., et al., Rhodopsin 5- and Rhodopsin 6-mediated clock synchronization in Drosophila melanogaster is independent of retinal phospholipase C-beta signaling. J Biol Rhythms, 2012. 27(1): p. 25–36.

30. Ni, J.D., et al., A rhodopsin in the brain functions in circadian photoentrainment in Drosophila. Nature, 2017. 545(7654): p. 340–344.

31. Au, D.D., et al., Mosquito cryptochromes expressed in Drosophila confer species-specific behavioral light responses. Curr Biol, 2022. 32(17): p. 3731–3744 e4.

32. Wan, G., et al., Cryptochrome 1 mediates light-dependent inclination magnetosensing in monarch butterflies. Nat Commun, 2021. 12(1): p. 771.

33. Froy, O., et al., Illuminating the circadian clock in monarch butterfly migration. Science, 2003. 300(5623): p. 1303–5.

34. Markert, M.J., et al., Genomic Access to Monarch Migration Using TALEN and CRISPR/Cas9-Mediated Targeted Mutagenesis. G3 (Bethesda), 2016. 6(4): p. 905–15.

35. Iiams, S.E., et al., Photoperiodic and clock regulation of the vitamin A pathway in the brain mediates seasonal responsiveness in the monarch butterfly. Proc Natl Acad Sci U S A, 2019.

36. Merlin, C., R.J. Gegear, and S.M. Reppert, Antennal circadian clocks coordinate sun compass orientation in migratory monarch butterflies. Science, 2009. 325(5948): p. 1700–4.

37. Rubin, E.B., et al., Molecular and phylogenetic analyses reveal mammalian-like clockwork in the honey bee (Apis mellifera) and shed new light on the molecular evolution of the circadian clock. Genome Res, 2006. 16(11): p. 1352–65.

38. Zhu, H., et al., Cryptochromes define a novel circadian clock mechanism in monarch butterflies that may underlie sun compass navigation. PLoS Biol, 2008. 6(1): p. e4.

39. Zhu, H., et al., The two CRYs of the butterfly. Curr Biol, 2005. 15(23): p. R953–4.

40. Helfrich-Forster, C., Light input pathways to the circadian clock of insects with an emphasis on the fruit fly Drosophila melanogaster. J Comp Physiol A Neuroethol Sens Neural Behav Physiol, 2020. 206(2): p. 259–272.

41. Sandrelli, F., et al., Comparative analysis of circadian clock genes in insects. Insect Mol Biol, 2008. 17(5): p. 447–63.

42. Currie, J., T. Goda, and H. Wijnen, Selective entrainment of the Drosophila circadian clock to daily gradients in environmental temperature. BMC Biol, 2009. 7: p. 49.

43. Wheeler, D.A., et al., Behavior in light-dark cycles of Drosophila mutants that are arrhythmic, blind, or both. J Biol Rhythms, 1993. 8(1): p. 67–94.

44. Brown, S.A., et al., Rhythms of mammalian body temperature can sustain peripheral circadian clocks. Curr Biol, 2002. 12(18): p. 1574–83.

45. Buhr, E.D., S.H. Yoo, and J.S. Takahashi, Temperature as a universal resetting cue for mammalian circadian oscillators. Science, 2010. 330(6002): p. 379–85.

46. Chen, C., et al., Drosophila Ionotropic Receptor 25a mediates circadian clock resetting by temperature. Nature, 2015. 527(7579): p. 516–20.

47. Chen, C., et al., nocte Is Required for Integrating Light and Temperature Inputs in Circadian Clock Neurons of Drosophila. Curr Biol, 2018. 28(10): p. 1595–1605 e3.

48. Malpel, S., A. Klarsfeld, and F. Rouyer, Larval optic nerve and adult extra-retinal photoreceptors sequentially associate with clock neurons during Drosophila brain development. Development, 2002. 129(6): p. 1443–53.

49. Park, P.S., Constitutively active rhodopsin and retinal disease. Adv Pharmacol, 2014. 70: p. 1–36.

50. Luo, D.G., et al., Apo-Opsin and Its Dark Constitutive Activity across Retinal Cone Subtypes. Curr Biol, 2020. 30(24): p. 4921–4931 e5.

51. Holliday, I.E. Two-Way ANOVA (v1.0.6) in Free Statistics Software (v1.2.1). 2019; Available from: https://www.wessa.net/rwasp_Two%20Factor%20ANOVA.wasp/.

52. Kirkman, T.W. Statistics to Use. 1996; Available from: http://www.physics.csbsju.edu/stats/.

53. Wu, G., et al., MetaCycle: an integrated R package to evaluate periodicity in large scale data. Bioinformatics, 2016. 32(21): p. 3351–3353.

